# Gene Expression Meta-Analysis of Colon Rectal Cancer Tumour Cells Reveals Genes in Association With Tumorogenesis

**DOI:** 10.1101/2022.01.30.478381

**Authors:** Rutvi Vaja

## Abstract

**Background:** Every year, more than 12 million people are diagnosed with colorectal cancer(CRC), and more than 600,000 people die from it, making it second most deadly form of cancer.This work analyzes differential gene expression across CRC and other glandular tumour samples to identify expression changes potentially contributing to the development of CRC tumorogenesis.

**Methods:** This work defines 13 gene signatures representing four CRC tumour and 10 other glandular tumours that are colonic by origin.Gene Set Enrichment Analysis (GSEA) is used to define positive and negative CRC gene panels from GSEA-identified leading-edge genes using two CRC signatures. GSEA then is used to verify enrichment and leading-edge gene membership of CRC panels in two independent CRC gene signatures. Analysis is then extended to four individual and 10 glandular tumour signatures. Genes most associated with CRC tumorogenesis are predicted by intersecting membership of GSEA-identified leading-edges across signatures.

**Results:** Significant enrichment is observed between CRC gene identification signatures, from which the positive (55 genes) and negative (77 genes) CRC panels are defined. Non-random significant enrichment is observed between CRC gene panels and verification signatures, from which 54 over- and 72 under-expressed genes are shared across leading-edges. Considering other glandular tumour samples individually and in combination with CRC, significant non-random enrichment is observed across these signatures. Eight solute carrier family genes such as (SLC25A32, SLC22A3, SLC25A20, SLC36A1, SLC26A3,SLC9A2, SLC4A4 and SLC26A2) from the CRC panel were shared commonly across all the gene signatures leading-edges, regardless of the colonic tumour type.

**Conclusion:** This meta-analysis identifies gene expression changes associated with the process of CRC tumorogenesis. These changes may contribute to developing therapeutic treatments available for CRC patients.

## 1. INTRODUCTION

As per the statistics of World Health Organization(WHO), Colorectal cancer (CRC) is the third most prevalent cancer in the world along with fourth leading cause of cancer related deaths(1).Colon and rectal cancers account for most of the glandular malignancies with the incidences increasing with age(1). Highly penetrant, autosomal dominantly or recessively inherited tumour predispositions cause about 5% of all colorectal cancers(2). More than 945 000 people are diagnosed with colorectal cancer each year, with roughly 492 000 patients dying(3). This type of cancer occurs infrequently, often as a result of genetic cancer syndromes or inflammatory bowel illnesses(3). As per the GLOBOCAN database documented 1.8 million newly diagnosed cases of CRC and 861,600 cases of CRC-related mortality over the world(4). CRC is a highly diverse disease caused mostly by interactions between genetic changes and environmental variables(5). Despite advances in diagnosis and treatment, the survival rate of CRC patients has remained unchanged over the previous two decades, with more than half of patients having regional or distant metastasis at the time of diagnosis(8).Several genes and cellular signalling pathways, including RACK1 (receptor for activated C kinase 1) and long noncoding RNA breast cancer anti-estrogen resistance 4 (lncRNA BCAR4), have been implicated in the formation and progression of CRC(6). RACK1 expression, for example, has been shown to be considerably upregulated in CRC tissues when compared to adjacent normal tissues(6). Despite these comprehensive and meticulous research(4–6) to find novel targets for CRC management, a comprehensive description of the critical key genes and signalling pathways implicated in CRC is lacking, to the best of our knowledge.

A high-throughput method for detecting mRNA expression in tissues, gene microarray profile analysis, is becoming a more promising tool in medical oncology. An enhanced understanding of the molecular pathogenesis of many cancer types can be gained by analysing differential gene expression between tumour tissues and normal control tissues, allowing for the identification of prospective target genes and signalling pathways for precision medicine(7). In earlier decades, microarray technology was employed in various research on gene expression profiles in cancer, but only one study focused on CRC(8). Apat from these studies, the comparative analysis of differentially expressed genes (DEGs) remains relatively limited(9). Furthermore, more research is needed to identify meaningful leading edge gene profiles for distinguishing CRC from normal tissues. In addition, the relationships among the DEGs should be clarified, along with the interaction networks and critical biological signalling pathways affected.

Many data mining analyses of mRNA, microRNA, long non-coding RNA, and DNA methylation have been performed on human cancers, particularly colon cancer, over the last few decades. A complete understanding about the molecular changes associated with CRC Tumor formation might help in assisting the development of new therapeutic interventions. Several studies have been done to elucidate molecular changes associated with CRC by examining gene expression changes in CRC-tumour forming cell cultures(10–12). There are some studies which provide a clinical reference for predicting the survival probability of patients with different clinical subtypes(13). Other studies geneerated protein-protein interaction (PPI) networks, and centrality analysis aswell was performed in order to identify the crucial genes that were potentially involved in the development of CRC(11). Several bioinformatics tools and strategies have aided in the exploration of molecular mechanisms of tumour pathogenesis and provided clues for better knowledge of related malignancies by identifying early biomarkers and potential therapeutic targets of tumours. Colon cancer is a multifactorial disease caused by a variety of factors including genetic, environmental, and lifestyle influences, but the pathogenesis of the disease is yet unknown (Aran et al., 2016). Exploring and analysing colon cancer’s molecular basis and important genes is critical to improve colon cancer prevention and treatment.

Previous work using a GSEA-based meta-analysis approach successfully identified known and novel genes associated with severe acute respiratory syndrome (SARS) infection through differential gene expression comparison between mRNA expression datasets (Park & Harris, 2021). Therefore, this paper applied the same GSEA-based approach to analyze mRNA expression data of tumor and normal CRC tissue derived from human colonic biopsy samples by defining and comparing gene expression signatures (i.e., gene lists ranked by differential expression)(15).

In this particular study, we have used expression profiling array datasets to identify and verify gene expression changes associated with CRC tumour formation to elucidate molecular changes associated with the process of tumorogenesis in CRC pathogenesis. Gene expression changes associated with CRC tumour formation then were compared to changes resulting from other colonic tumour forming diseases like Ulcerative coilitis, adenoma dyslapsia and Hyperplastic Polyp, to examine the common leading edge genes in CRC that invoked changes and played a potential role in CRC tumour formatio. Finally we verified the gene signatures in several other colonic tumour forming diseases such as Ulcerative coilitis, adenoma dyslapsia and Hyperplastic Polyp. A meta-analysis was performed lastly to find the common shared leading edge genes across all different conditions of CRC. The gene expression changes identified with across these several conditions holds the potential to actually improve the treatments targetted for CRC and it’s therpeutic intervention.

## 2. METHODS

### 2.1 mRNA Expression Resources

To identify gene expression changes associated with CRC tumour forming cells, the Gene Expression Omnibus (GEO) repository (16) was searched to find datasets for use in this study (Table 1). The six independent data series GSE44861,GSE113513, GSE44076 GSE10714,GSE32323 and GSE24514 had CRC tumour and normal tissue samples and hence we started our analysis from here. GSE44861 is an Affymetrix expression data collected from colon cancer patient tissues in which RNA from fresh frozen colon tissues were extracted using Trizol and hybridized to Affymetrix U113A arrays(17). The platform used for GSE44861 was GPL3921 [HT_HG-U133A] Affymetrix HT Human Genome U133A Array. GSE113513 has samples from 14 colorectal cancer patients who had undergone surgical resection of colorectal cancer where Trizol (Thermo Fisher Scientific) extraction of total RNA was performed(18). The platform used for GSE113513 was GPL15207 [PrimeView] Affymetrix Human Gene Expression Array. GSE44076 contained Gene expression profiles of paired normal adjacent mucosa and tumor samples from 98 individuals and 50 healthy colon mucosae, which were obtained through Affymetrix Human Genome U219 Arrays(19). The platform used for GSE44076 was GPL13667 [HG-U219] Affymetrix Human Genome U219 Array. GSE10714 had expression data from human colonic biopsy samples on which Qiagen RNeasy Mini extraction of total RNA was performed(20). The platformused for GSE10714 was GPL570 [HG-U133_Plus_2] Affymetrix Human Genome U133 Plus 2.0 Array. GSE32323 consisted of gene expression profiles for 17 pairs of cancer and non-cancerous tissues from colorectal cancer patients were measured by Affymetrix HG-U133 Plus 2.0 arrays. Here, total RNA was extracted from tissue specimens using RNeasy kit (Qiagen, Hilden, Germany)(21).The platform used for GSE32323 was GPL570 [HG-U133_Plus_2] Affymetrix Human Genome U133 Plus 2.0 Array. Lastly, GSE24514 had expression data from human MSI colorectal cancer and normal colonic mucosa in which RNA rom fresh frozen tissues was extracted with Trizol reagent (Invitrogen)(22). The platform used for GSE24514 was GPL96 [HG-U133A] Affymetrix Human Genome U133A Array].

**TABLE 1.** Datasets Utilized for this Study.

| Data-Set | Description | Platform | Probes | Genes |
| --- | --- | --- | --- | --- |
| GSE44861 | Gene expression profiling of 111 colon tissues from tumors and adjacent noncancerous tissues. RNA from fresh frozen colon tissues were extracted using Trizol and hybridized to Affymetrix U113A arrays | GPL3927 | 22277 | 21248 |
| GSE113513 | Samples from fourteen colorectal cancer patients who had undergone surgical resection of colorectal cancer. Trizol (Thermo Fisher Scientific) extraction of total RNA was performed on the samples | GPL15207 | 49395 | 48872 |
| GSE44076 | Paired normal adjacent mucosa and tumor samples from 98 individuals and 50 healthy colon mucosae. | GPL13667 | 49386 | 48785 |
| GSE10714 | Expression data from human colonic biopsy sample.Total RNA was extracted from colonic biosy samples CRC and hybridized on Affymetrix HGU133 Plus 2.0 microarrays | GPL570 | 54675 | 45782 |
| GSE32323 | Gene expression profiles for 17 pairs of cancer and non-cancerous tissues from colorectal cancer patients were measured by Affymetrix HG-U133 Plus 2.0 arrays. | GPL570 | 54675 | 45782 |
| GSE24514 | Expression profiling of 34 MSI colorectal cancers and 15 normal colonic mucosas . Samples had Comparison of malignant and healthy tissue. | GPL96 | 22283 | 21225 |

The expression data provided by GEO for all datasets were z-scored normalized across all samples within the dataset regardless of tumour cell or treatment type prior to use for analysis. The expression data was cleaned by removing probe identifiers given by GEO where 1) all samples having gene expression z-score of 0, or 2) duplicate identifiers/symbols were identified so only the identifier with the highest coefficient of variation were retained. The soft family data consisted of Entrez Gene ids, Gene symbols and Probe identifiers as well. If a dataset’s GEO-provided platform contained both Ensemble gene IDs and gene symbols for a probe, the GEO-provided platform files to convert between these two prove identifiers were used. For this particular study, we have converted the probe identifiers to gene symbols using the Database for Annotation, Visualization, and Integrated Discovery (DAVID) v6.8 gene conversion tool(23) in order to maintain uniformity.In order to obtain other metadata information like sample characteristics, sample source, Ensemble ids and GSM numbers, GEO-provided platform files were used.

### 2.2 Defining Gene Signatures

In order to examine the gene expression changes associated with umour formation in CRC differential gene expression was measured for samples of interest from each dataset using Welch’s two-sample Ttest score of cleaned and normalized values. (For eg: tumour samples and normal tissue sample from one dataset formed one signature). Samples with the same origin, regardless of tumour stage, cancer type(metastasis or beningn),were combined to form one signature. The resulting list of genes along with their T-test scores were used to define 13 gene signatures. This signatures are formed from the gene lists ranked from high to low differential gene expression between tumor versus normal tissue samples (Park & Harris, 2021). The signatures derived from the same dataset used the same control samples. The gene location where T-score becomes negative (i.e., T-score=0) and the T-score range for each signature are found in Table 2.

**TABLE 2.**
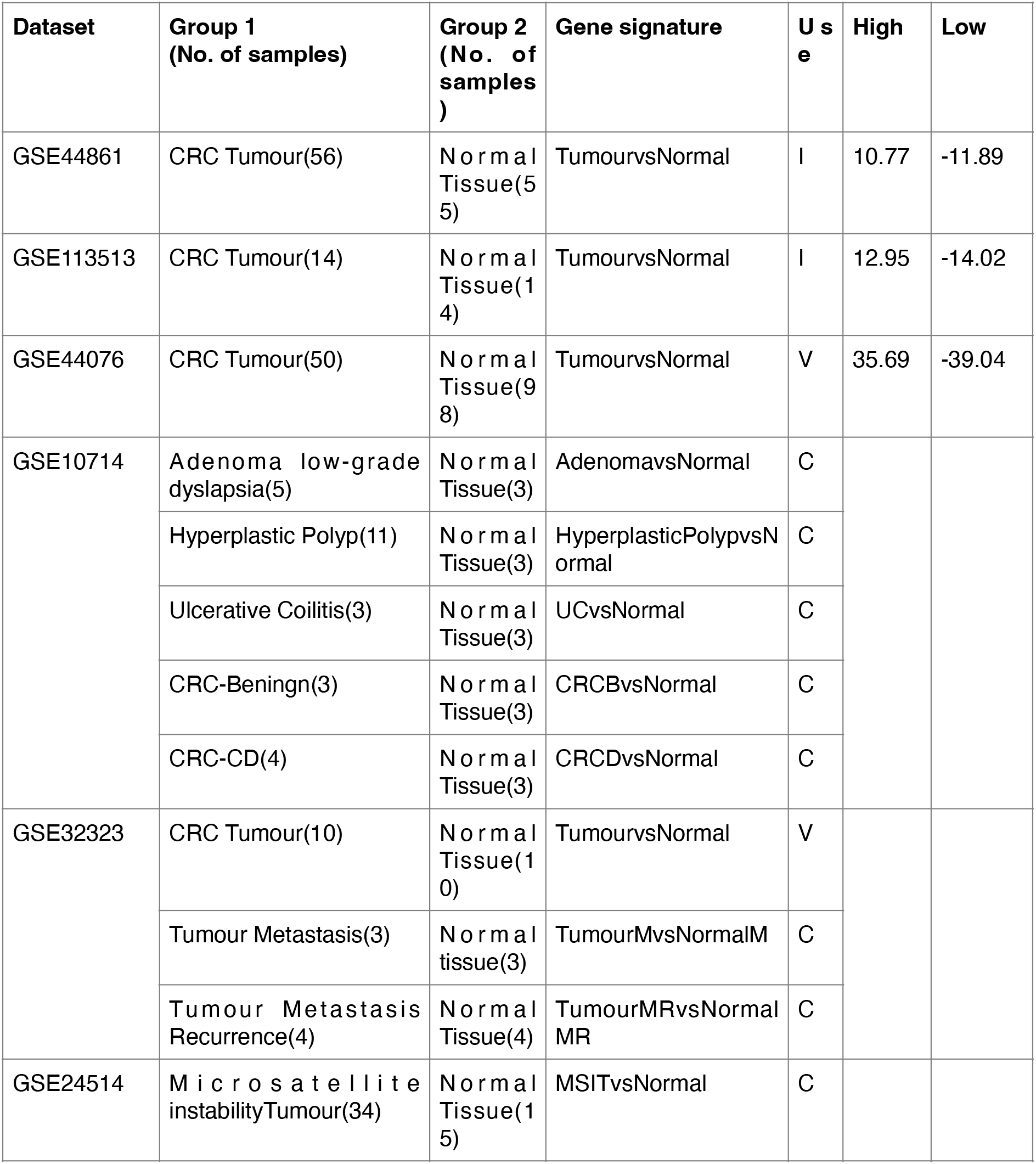
Signatures Defined in this Study.

| Dataset | Group 1<br>(No. of samples) | Group 2<br>(No. of samples) | Gene signature | Use | High | Low |
| --- | --- | --- | --- | --- | --- | --- |
| GSE44861 | CRC Tumour(56) | Normal Tissue(55) | TumourvsNormal | I | 10.77 | -11.89 |
| GSE113513 | CRC Tumour(14) | Normal Tissue(14) | TumourvsNormal | I | 12.95 | -14.02 |
| GSE44076 | CRC Tumour(50) | Normal Tissue(98) | TumourvsNormal | V | 35.69 | -39.04 |
| GSE10714 | Adenoma low-grade dyslapsia(5) | Normal Tissue(3) | AdenomavsNormal | C |  |  |
|  | Hyperplastic Polyp(11) | Normal Tissue(3) | HyperplasticPolypvsNormal | C |  |  |
|  | Ulcerative Coilitis(3) | Normal Tissue(3) | UCvsNormal | C |  |  |
|  | CRC-Beningn(3) | Normal Tissue(3) | CRCBvsNormal | C |  |  |
|  | CRC-CD(4) | Normal Tissue(3) | CRCDvsNormal | C |  |  |
| GSE32323 | CRC Tumour(10) | Normal Tissue(10) | TumourvsNormal | V |  |  |
|  | Tumour Metastasis(3) | Normal tissue(3) | TumourMvsNormalM | C |  |  |
|  | Tumour Metastasis Recurrence(4) | Normal Tissue(4) | TumourMRvsNormalMR | C |  |  |
| GSE24514 | Microsatellite instabilityTumour(34) | Normal Tissue(15) | MSITvsNormal | C |  |  |

**TABLE 3.** Positive CRC Gene Panel Defined in this Study.

| Entrez ID | Gene Symbol | Description |
| --- | --- | --- |
| 9603 | NFE2L3 | nuclear factor (erythroid-derived 2)-like 3 |
| 7004 | TEAD4 | TEA domain family member 4 |
| 81034 | SLC25A32 | solute carrier family 25, member 32 /// solute carrier family 25, member 32 |
| 9569 | GTF2IRD1 | GTF2I repeat domain containing 1 |
| 1876 | E2F6 | E2F transcription factor 6 |
| 57460 | PPM1H | protein phosphatase 1H (PP2C domain containing) |
| 8886 | DDX18 | DEAD (Asp-Glu-Ala-Asp) box polypeptide 18 |
| 5471 | PPAT | phosphoribosyl pyrophosphate amidotransferase |
| 6581 | SLC22A3 | solute carrier family 22 (extraneuronal monoamine transporter), member 3 |
| 11260 | XPOT | exportin, tRNA (nuclear export receptor for tRNAs) |
| 54517 | PUS7 | pseudouridylate synthase 7 homolog (S. cerevisiae) |
| 54529 | ASNSD1 | asparagine synthetase domain containing 1 |
| 1312 | COMT | catechol-O-methyltransferase |
| 595 | CCND1 | cyclin D1 |
| 60496 | AASDHPPT | aminoadipate-semialdehyde dehydrogenase-phosphopantetheinyl transferase |
| 1662 | DDX10 | DEAD (Asp-Glu-Ala-Asp) box polypeptide 10 |
| 6059 | ABCE1 | ATP-binding cassette, sub-family E (OABP), member 1 |
| 10807 | SDCCAG3 | serologically defined colon cancer antigen 3 |
| 52 | ACP1 | acid phosphatase 1, soluble |
| 55795 | PCID2 | PCI domain containing 2 |
| 10196 | PRMT3 | protein arginine methyltransferase 3 |
| 63875 | MRPL17 | mitochondrial ribosomal protein L17 |
| 79728 | PALB2 | partner and localizer of BRCA2 |
| 1615 | DARS | aspartyl-tRNA synthetase |
| 5361 | PLXNA1 | plexin A1 |
| 26586 | CKAP2 | cytoskeleton associated protein 2 |
| 79074 | C2orf49 | chromosome 2 open reading frame 49 |
| 57122 | NUP107 | nucleoporin 107kDa |
| 8609 | KLF7 | Kruppel-like factor 7 (ubiquitous) |
| 11146 | GLMN | glomulin, FKBP associated protein |
| 2118 | ETV4 | ets variant gene 4 (E1A enhancer binding protein, E1AF) /// ets variant gene 4 (E1A enhancer binding protein, E1AF) |
| 6282 | S100A11 | S100 calcium binding protein A11 |
| 23560 | GTPBP4 | GTP binding protein 4 |
| 8833 | GMPS | guanine monphosphate synthetase |
| 10171 | RCL1 | RNA terminal phosphate cyclase-like 1 |
| 3614 | IMPDH1 | IMP (inosine monophosphate) dehydrogenase 1 |
| 54881 | TEX10 | testis expressed sequence 10 |
| 26009 | ZZZ3 | zinc finger, ZZ-type containing 3 |
| 26031 | OSBPL3 | oxysterol binding protein-like 3 |
| 51776 | ZAK | sterile alpha motif and leucine zipper containing kinase AZK |
| 26135 | SERBP1 | SERPINE1 mRNA binding protein 1 |
| 4233 | MET | met proto-oncogene (hepatocyte growth factor receptor) |
| 79084 | WDR77 | WD repeat domain 77 |
| 9128 | PRPF4 | PRP4 pre-mRNA processing factor 4 homolog (yeast) |
| 10527 | IPO7 | importin 7 |
| 6741 | SSB | Sjogren syndrome antigen B (autoantigen La) |
| 22880 | MORC2 | MORC family CW-type zinc finger 2 |
| 9221 | NOLC1 | nucleolar and coiled-body phosphoprotein 1 |
| 1736 | DKC1 | dyskeratosis congenita 1, dyskerin |

**TABLE 4.** Negative CRC Gene Panel Defined in this Study

| Entrez Id | Gene symbol | Description |
| --- | --- | --- |
| 25840 | METTL7A | methyltransferase like 7A |
| 51196 | PLCE1 | phospholipase C, epsilon 1 |
| 11099 | PTPN21 | protein tyrosine phosphatase, non-receptor type 21 |
| 1908 | EDN3 | endothelin 3 |
| 23228 | PLCL2 | phospholipase C-like 2 |
| 51228 | GLTP | glycolipid transfer protein |
| 2110 | ETFDH | electron-transferring-flavoprotein dehydrogenase |
| 55743 | CHFR | checkpoint with forkhead and ring finger domains |
| 10223 | GPA33 | glycoprotein A33 (transmembrane) |
| 788 | SLC25A20 | solute carrier family 25 (carnitine/acylcarnitine translocase), member 20 |
| 54884 | RETSAT | retinol saturase (all-trans-retinol 13,14-reductase) |
| 5873 | RAB27A | RAB27A, member RAS oncogene family |
| 171586 | ABHD3 | abhydrolase domain containing 3 |
| 55359 | STYK1 | serine/threonine/tyrosine kinase 1 |
| 54474 | ALOX | keratin 20 |
| 6414 | SEPP1 | selenoprotein P, plasma, 1 |
| 3960 | LGALS4 | lectin, galactoside-binding, soluble, 4 (galectin 4) |
| 51411 | BIN2 | bridging integrator 2 |
| 140803 | TRPM6 | transient receptor potential cation channel, subfamily M, member 6 |
| 125 | ADH1B | alcohol dehydrogenase IB (class I), beta polypeptide |
| 80031 | SEMA6D | sema domain, transmembrane domain (TM), and cytoplasmic domain, (semaphorin) 6D |
| 5567 | PRKACB | protein kinase, cAMP-dependent, catalytic, beta |
| 5827 | PXMP2 | peroxisomal membrane protein 2, 22kDa |
| 64922 | LRRC19 | leucine rich repeat containing 19 |
| 2168 | FABP1 | fatty acid binding protein 1, liver |
| 9218 | VAPA | VAMP (vesicle-associated membrane protein)-associated protein A, 33kDa |
| 23171 | GPD1L | glycerol-3-phosphate dehydrogenase 1-like |
| 55620 | STAP2 | signal-transducing adaptor protein-2 |
| 7263 | TST | thiosulfate sulfurtransferase (rhodanese) |
| 206358 | SLC36A1 | solute carrier family 36 (proton/amino acid symporter), member 1 |
| 4128 | MAOA | monoamine oxidase A |
| 8857 | FCGBP | Fc fragment of IgG binding protein |
| 51316 | PLAC8 | placenta-specific 8 |
| 10351 | ABCA8 | ATP-binding cassette, sub-family A (ABC1), member 8 |
| 9060 | PAPSS2 | 3'-phosphoadenosine 5'-phosphosulfate synthase 2 |
| 10891 | PPARGC1A | peroxisome proliferator-activated receptor gamma, coactivator 1 alpha |
| 55286 | C4orf19 | chromosome 4 open reading frame 19 |
| 25994 | HIGD1A | HIG1 domain family, member 1A |
| 10924 | SMPDL3A | sphingomyelin phosphodiesterase, acid-like 3A |
| 1087 | CEACAM7 | carcinoembryonic antigen-related cell adhesion molecule 7 |
| 2517 | FUCA1 | fucosidase, alpha-L- 1, tissue |
| 2647 | BLOC1S1 | biogenesis of lysosome-related organelles complex-1, subunit 1 |
| 771 | CA12 | carbonic anhydrase XII |
| 58472 | SQRDL | sulfide quinone reductase-like (yeast) |
| 760 | CA2 | carbonic anhydrase II |
| 3957 | LGALS2 | lectin, galactoside-binding, soluble, 2 (galectin 2) /// lectin, galactoside-binding, soluble, 2 (galectin 2) |
| 7102 | TSPAN7 | tetraspanin 7 |
| 11148 | HHLA2 | HERV-H LTR-associating 2 |
| 4306 | NR3C2 | nuclear receptor subfamily 3, group C, member 2 |
| 9314 | KLF4 | Kruppel-like factor 4 (gut) |
| 10050 | SLC17A4 | solute carrier family 17 (sodium phosphate), member 4 |
| 81618 | ITM2C | integral membrane protein 2C /// integral membrane protein 2C |
| 2494 | NR5A2 | nuclear receptor subfamily 5, group A, member 2 |
| 35 | ACADS | acyl-Coenzyme A dehydrogenase, C-2 to C-3 short chain |
| 1811 | SLC26A3 | solute carrier family 26, member 3 |
| 6549 | SLC9A2 | solute carrier family 9 (sodium/hydrogen exchanger), member 2 |
| 957 | ENTPD5 | ectonucleoside triphosphate diphosphohydrolase 5 |
| 2767 | GNA11 | guanine nucleotide binding protein (G protein), alpha 11 (Gq class) |
| 608 | TNFRSF17 | tumor necrosis factor receptor superfamily, member 17 |
| 10917 | BTNL3 | butyrophilin-like 3 |
| 3291 | HSD11B2 | hydroxysteroid (11-beta) dehydrogenase 2 |
| 7358 | UGDH | UDP-glucose dehydrogenase |
| 8671 | SLC4A4 | solute carrier family 4, sodium bicarbonate cotransporter, member 4 |
| 123887 | ZG16 | zymogen granule protein 16 |
| 5794 | PTPRH | protein tyrosine phosphatase, receptor type, H |
| 5333 | PLCD1 | phospholipase C, delta 1 |
| 79148 | MMP28 | matrix metalloproteinase 28 |
| 10590 | SCGN | secretogogin, EF-hand calcium binding protein |
| 23140 | ZZEF1 | zinc finger, ZZ-type with EF-hand domain 1 |
| 1836 | SLC26A2 | solute carrier family 26 (sulfate transporter), member 2 |
| 11240 | PADI2 | peptidyl arginine deiminase, type II |
| 27250 | PDCD4 | programmed cell death 4 (neoplastic transformation inhibitor) |
| 762 | CA4 | carbonic anhydrase IV |
| 759 | CA1 | carbonic anhydrase I |
| 1113 | CHGA | chromogranin A (parathyroid secretory protein 1) |
| 2980 | GUCA2A | guanylate cyclase activator 2A (guanylin) |

### 2.3 Identification of Genes Associated with CRC Tumour Tissues

To identify gene expression changes associated with CRC Tumour tissues, two CRC tumour gene panels were generated (Figure 1). In order to do this, 500 genes from the positive and negative tails from both the GSE44861-derived TumourvsNormal and GSE113513-derived TumourvsNormal gene signatures were selected and then were used to form four individual query gene sets. GSEA compared each query gene set to the both the entire GSE44861-derived TumourvsNormal and GSE113513-derived TumourvsNormal gene signatures(reference). For our identification stage, we used the datasets GSE44861 as query first against GSE113513 as reference, followed by GSE113513 as query dataset against the whole of GSE44861 TumourvsNormal signature.Leading-edge (LE) genes from each of these analysis were examined and shared leading-edge genes were used to define two CRC TumourvsNormal gene panels, one panel per tail(Positive CRC and Negative CRC Panels). Pathway enrichment analysis was performed on both CRC tumour gene panels using DAVID.

**FIGURE 1:**
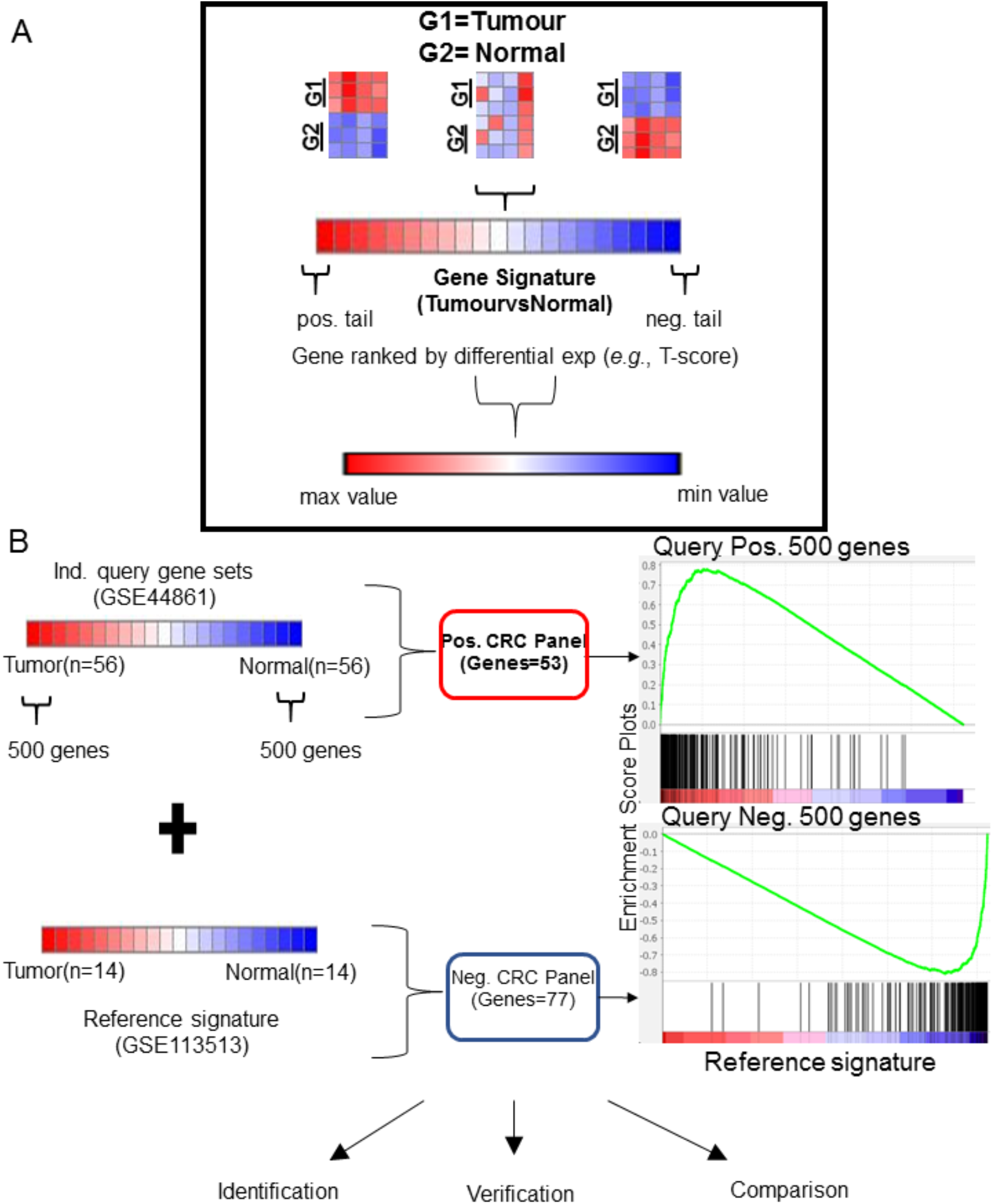
Schematic Overview of Gene Panel Identification.

### 2.4 Verification of CRC Tumour Gene Panels

To verify the CRC Tumour gene panels, GSEA between CRC Tumour gene panels(Pos. CRC and Neg. CRC) and GSE44076-derived and GSE10714-derived TumourvsNormal signatures was performed. To assess if results generated from GSEA could be achieved randomly, 1000 gene panels consisting of 175-genes to match the average number of genes in the positive and negative CRC Tumour panels(query signature) were randomly selected from the GPL3927 platform used to define the GSE44861 gene signatures used for gene identification for GSEA against GSE44076-derived and GSE32323-derived TumourvsNormal signatures (reference signature). These analysis generated a null distribution of NES(Normalized enrichment score) to which were compared the NES achieved by CRC tumour gene panels for each reference gene signature and count the number of equal or better NES to estimate significance (i.e., null distribution p-value). The bar and whiskers Plot was created and calculated using Excel. Heatmaps were generated by a user-friendly, web-based program maps.https://software.broadinstitute.org/morpheus. Morpheus is a softwrae created by Broad Institute Software.

### 2.5 Comparison of CRC Tumour Gene Panels to Other types of tumour that are Glandular in Origin

GSEA was used to compare the identified CRC gene panels and gene signatures derived from samples of one of the six glandular tumour (Ulcerative Coilitis, Hyperplastic Polyp, Adenoma with low grade dylapsia, Colorectal cancer –Beningn,Colorectal cancer with a chronic disease and colorectal Tumours with microsatellite instability), compared to normal tissue samples was performed to compare gene expression changes across CRC tumour tissues and several other glandular tumours. Following this, random modelling as stated earlier was used to assess if results generated from GSEA could be achieved randomly. Leading-edge genes from each statistically significant (GSEA p-value<0.05), non-random (null distribution p-value<0.05) GSEA were examined and analysed for common genes.

## 3. RESULTS

### 3.1 Gene Signature Approach Identified Gene Expression Changes Asspciated with Colorectal Cancer Tumor Tissues *in vitro*

GSE44861-derived and GSE113513-derived TumourvsNormal gene signatures were defined to identify genes associated with response to Tumour tissue of human CRC colonic biopsy samples. (Table 2). From both these two TumourvsNormal identification signatures, two gene sets were generated containing the 500 most differentially expressed genes from the positive and negative tails of each signature, capturing maximum coverage of the signature that was allowable by GSEA(24). The T-score values for GSE44861 derived TumourvsNormal signature was >10.77 and <-11.89. For GSE113513 derived TumourvsNormal gene signature was >12.95 and <-14.0282. To find the similarity between these two signatures, enrichment was first calculated using GSEA between GSE113513-derived TumourvsNormal positive or negative tail gene sets (individual query set) and the GSE44861-derived TumourvsNormal(reference set) and achieved NES=3.54 and NES=-3.64 for positive and negative tail query gene sets, respectively, both with a GSEA p-value<0.001. Similarly enrichment was calculated using GSEA between GSE44861-derived TumourvsNormal positive or negative tail gene sets (individual query set) and the GSE113513-derived TumourvsNormal (reference set) and achieved NES=3.38 and NES=-4.14 for positive and negative tail query gene sets, respectively, both with a GSEA p-value<0.001. The identified leading-edge genes from GSEA are listed in Supplemental Material STables 1.

No genes in the positive CRC panel were mentioned in the published reports for GSE44861 and GSE113513, though some panel genes like Kruppel like factor-7(ubiquitious) (KLF-7) and RNA terminal phosphate cyclase-like 1(RCL-1) have reported connections with CRC tumour formation in the literature (27,28). In the negative CRC gene panel, Keratin 20(ALOX) was found to have previous associations with CRC tumour formation(29). Taking all the results together, this demonstrated the detection ability of using a GSEA-based approach to gene identification. The rest of the genes in the negative panobinostat panel had no prior association with CRC tumour formation from the published reports for GSE44861 and GSE113513. It can be speculated that genes lacking previously reported associations with CRC tumour formation that were identified here also are associated with human colonic biopsy samples.

To expand our analysis, the cellular roles of CRC gene panels were examined using DAVID to calculate enrichment between each CRC gene panel and pathways in popular known knowledgebases. It was noticed that, when compared to other databases, the GO BP database returned the most significantly enriched pathways (data not mentioned here), hence this discussion was concentrated on GO-BP data to prevent confusion caused by other overlapping pathway and gene inclusion differences across other multiple known knowledgebases.

DAVID identified nine significant GO-BP pathways (EASE score p-value<0.05) from the positive CRC gene panel and 10 significant pathways from the negative CRC gene panel (Supplemental Material STable 2). Some significantly enriched pathways have experimentally established associations with CRC tumour formation, such as RNA processing pathway (GO:0006396, p-value=0.02), demonstrating the ability of our gene signature approach to detect pathways associated with CRC Tumour formation(25). Other identified pathways, such as lipid catabolic process (GO:0016042, p-value=0.04), have no prior associations to CRC Tumour formation. Therefore, it can be speculated that pathways that came out as a result without prior association to CRC Tumours identified here also were involved in CRC.

### 3.2 Enrichment of CRC Gene Panels and Specific CRC Panel Genes Verified in Independent Datasets

To verify our CRC gene panels, GSEA was used to calculate enrichment between our CRC panels (individual queries) and two verification gene signatures (individual references): GSE32323-derived TumourvsNormal and GSE44076-derived TumourvsNormal (Table 2). Significant similarity between positive and negative CRC panels and GSE32323-derived TumourvsNormal (NES=2.26 for the positive CRC panel, Figure 2A, and NES=-2.56 for the negative CRC panel, Figure 2B, both GSEA p-value<0.001) was found. To determine how likely the NES achieved for CRC gene panels would be achieved by random chance, 1000 randomly selected 175-gene panels were generated from the GSE44861-derived TumourvsNormal gene signature to match the average size and potential composition of our CRC gene panels. GSEA was then repeated using these randomly generated gene panels (individual queries) and the GSE44076-derived TumourvsNormal (reference) to generate a null distribution of NES achieved via a random chance.

**FIGURE 2:**
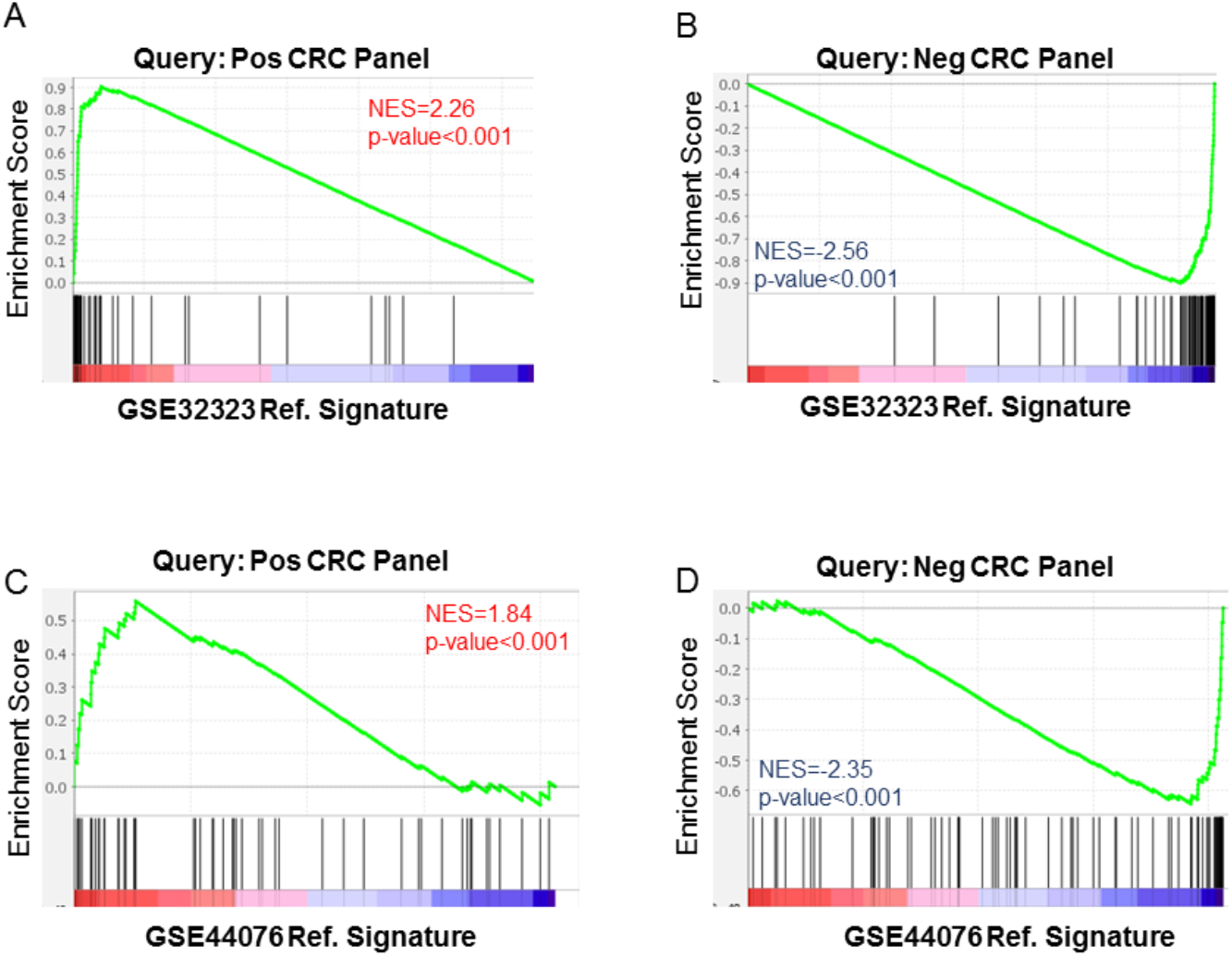
Schematic Overview of Gene Panel Identification

**FIGURE 3:**
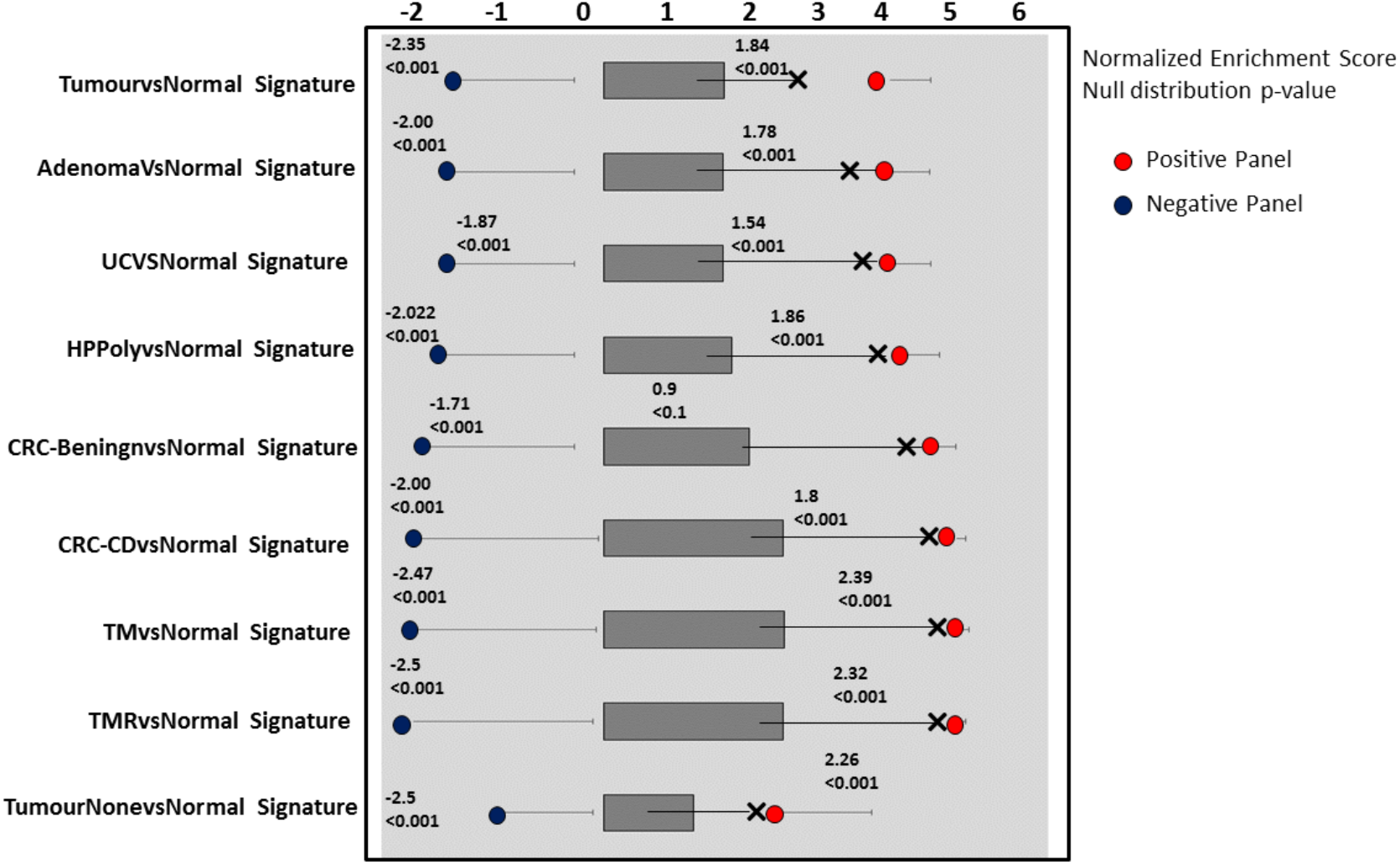
Enrichment of CRC Gene Panels and Random Models Across Other glandular Tumours

From this, random NES ranged from 1.47 to −1.5 was found (data not shown), illustrating that NES achieved by our CRC panels are non-random (null distribution p-value<0.001).Taken together, these results demonstrate that the enrichment achieved from our panobinostat panels was true.

To determine which of our CRC panel genes were verified across all signatures, leading-edge genes identified by GSEA for each verification signature were examined. Leading-edge genes for GSE44076-derived and GSE32323-derived TumourvsNormal signatures are listed in Supplemental Material STables 3 and 9, respectively. 51 genes from the positive CRC panel and 75 genes from the negative CRC panel were shared between verification signatures. These data together verify our shared leading-edge genes are associated with CRC tumour tissues in human colonic biopsy samples and support the hypothesis that identified genes without previously reported associations are also associated with CRC tumour tissues in human colonic biopsy samples.

### 3.3 Non-random Enrichment of CRC Tumour Gene Panels Found in Other Glandular Tumours

To expand this study, gene expression changes associated with CRC Tumour were compared to changes observed in other glandular tumours included in GEO Series previously used to examine the common shared leading edge genes between CRC tumour tissues and other glandular tumours. The following eight signatures were examined: AdenomavsNormal, HyperplasticPolypvsNormal, UcvsNormal, CRCBvsNormal, CRCDvsNormal, TumourMvsNormalM, TumourMRvsNormalMR and MSITvsNormal(Table 2). For AdenomavsNormal gene signature versus the positive panel NES=1.785 and for Negative panel is NES=-2.00(GSEA p. value<0.001). For HyperplasticPolyvsNormal gene signature, when running a query against the Positive panel NES=1.86 and for negative panel NES=-2.022(GSEA p. value<0.001). For UcvsNormal gene signature, NES for positive panel=1.54 and NES for negative panel is −1.87(GSEA p. value<0.001). For CRCBvsNormal gene signature against positive panel query, NES=0.9 and for negative panel query, NES=-1.7(GSEA p. value<0.001).For CRCDvsNormal gene signature, the query against Positive panel yielded a NES of 1.866 whereas the NES of negative panel was −2.00(GSEA p. value<0.001).The TumourMvsNormalM gene signature yielded NES of 2.39 for the positive gene panel as the query set and NES of −2.47(GSEA p. value<0.001).For TumourMRvsNormalMR the NES for the positive panel was 2.33 and for the negative panel NES=-2.5(GSEA p. value<0.001). For gene signature of MSITvsNormal the positive panel’s NES score is 2.307 and negative panel’s NES score is −3.12(GSEA p. value<0.001). Both the positive and negative CRC panel, showed significant enrichment score(GSEA p. value<0.001) in all the gene signatures with and were also non-random(null distribution p-values<0.001).

### 3.4 Leading-edge Genes Found Amongst the Other Glandular Tumours

Finally, leading-edge genes from each GSEA across other glandular tumours were examined to identify genes of potential interest. Supplemental Material STables 3 through 9 contains leading-edge genes from GSEA with each CRC gene panel across 8 comparison gene signatures (Table 2). Out of the 54 common shared genes in the Positive CRC panel across all the signatures, 48 genes stood out having a statistical significance of (p value<0.001). Whereas out of the 75 common shared genes in the negative CRC panel across all the signatures, 72 genes were of highly statistical significance of (p value<0.001).9 solute carrier family genes solute carrier family 25, member 32 /// solute carrier family 25 member 32(SLC25A32), solute carrier family 22 (extraneuronal monoamine transporter) member 3(SLC22A3), solute carrier family 25(carnitine/acylcarnitine translocase) member 20(SLC25A20),solute carrier family 36 (proton/amino acid symporter)member 1(SLC36A1),solute carrier family 17 (sodium phosphate)member 4(SLC17A4),solute carrier family 26 member 3(SLC26A3), solute carrier family 9 (sodium/hydrogen exchanger)member 2(SLC9A2), solute carrier family 4 sodium bicarbonate cotransporter member 4(SLC4A4),and solute carrier family 26 (sulfate transporter), member 2(SLC26A2) from the CRC panel were shared commonly across all the gene signatures. Group of carbonic anhydrase protein coding genes from CRC gene Panel were also commonly shared across all gene signature’s namely carbonic anhydrase XII(CA12), carbonic anhydrase II(CA2), carbonic anhydrase IV(CA4) and carbonic anhydrase I(CA1).

To rank shared leading-edge genes, T-test p value analysis using Excel-STAT software was used. Out of the 129 Leading edge genes(54 from positive gene panel and 75 from negative gene panel) we wanted to find out the significant leading edge genes. Hence using the parameter: p. Value<0.001 we applied a P-value T-test in Excel to calculate the P-value. A total of 120 genes(48-Over-expressed and 72 under-expressed) out of the 129 were significantly valid lleading edge genes. The volcano plot for the significant leading-edge genes is shown in Figure 4.

**FIGURE 4:**
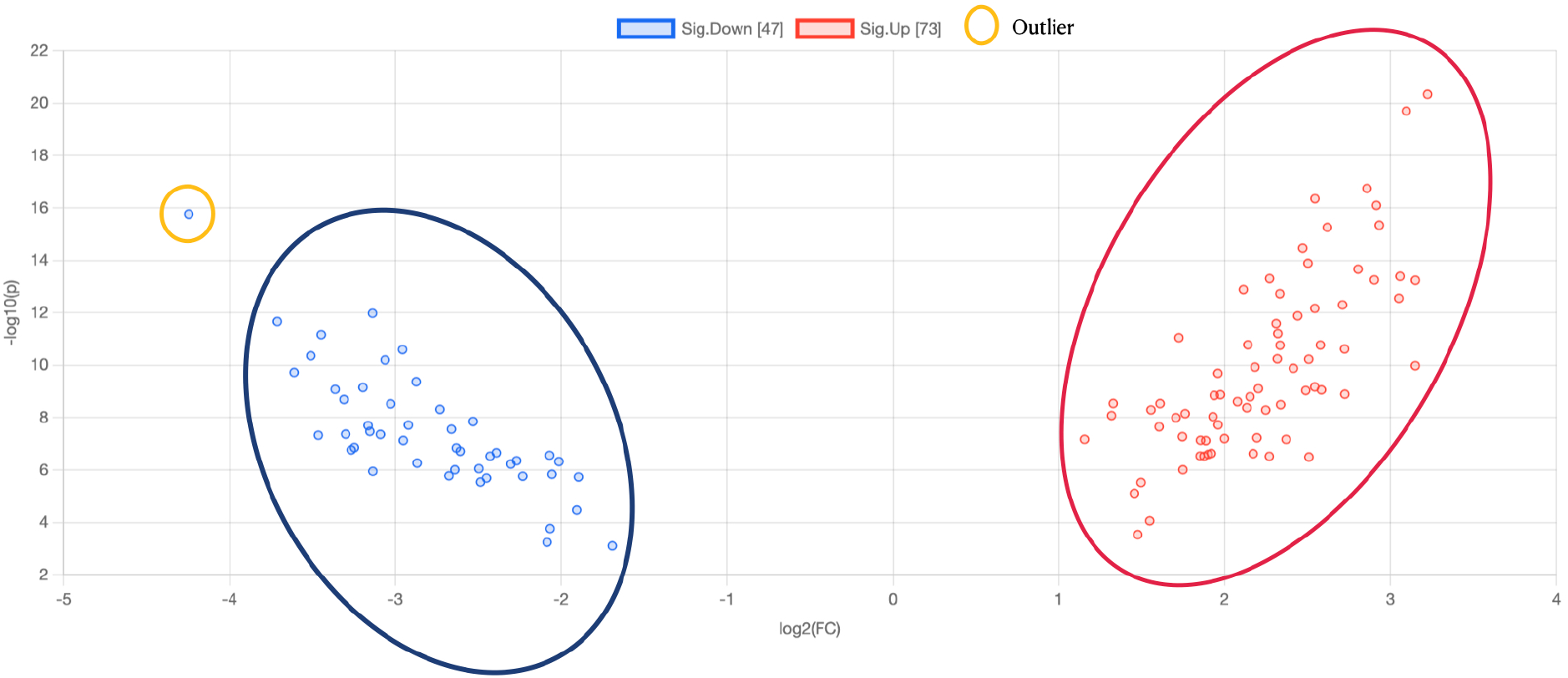
Volcano Plot showing over-expressed and Under-expressed leading edge Genes

## 4. DISCUSSION

Colorectal cancer still stands as a predominant cancer and is the second- and third-most common cancer in women and men, respectively(26). Despite considerable breakthroughs in treatment, colorectal cancer mortality remains high, with 40–50 percent of patients dying as a result of their illness. Identification of differentially expressed genes associated with CRC itself can contribute to the overall understanding of the molecular changes that drive cellular and molecular changes in CRC tumour development. This improved understanding can potentially contribute to the development of new therapeutic options to improve the prognosis for CRC patients. This work conducted a metaanalysis of gene expression signatures generated from mRNA expression data across CRC tumours and eight other glandular tumours which are colonic in nature,to identify differentially expressed genes associated with CRC tumour formation.Genes that change in response to treatment may contribute to developing treatment resistance long-term.

In this study, out of the 55 genes from the positive CRC gene panel, 54 of them were continously constant throughout all the gene signatures and for the negative gene CRC panel which had 77 genes, 75 of them were uniform across all the other gene signatures. To find out wether, these genes are statistically significant or not we did a T-test for calculating the p value in excel.Among the genes identified in this study, from the positive CRC gene panel, 54 common shared genes in the Positive CRC panel across all the signatures, 48 genes stood out having a statistical significance of (p value<0.001). Whereas out of the 75 common shared genes in the negative CRC panel across all the signatures, 72 genes were of highly statistical significance of (p value<0.001). Among the genes identified, studies have shown that nuclear factor (erythroid-derived 2)-like 3(NFE2L3) decreases colon cancer cell proliferation in vitro and tumor growth in vivo(30). Intrestingly, sterile alpha motif and leucine zipper containing kinase AZK(ZAK) gene has involved funcionalities in lung cancer tumorogenesis process and JNK pathway activation(31).Methyltransferase like 7A(METTL7A) have shown previous associations with thyroid cancer but not in lung, uterine, ovarian, gastric, esophagus, pancreatic, liver, or colorectal cancers via bioinformatic analysis(32).aken together, these results suggested that the GSEA-based meta-analysis approach used here was successful in identifying cancer-related genes with and without CRC tumorogenesis associations.

This work had observable gene detection limitations that may have biological implications. For example, gene expression changes commonly associated with CRC tumorogenesis in genes like MSH2 and MSH6 both on chromosome 2 and MLH1, on chromosome 3 were not found in this study (33,4,30). Platform variations, both in gene inclusion and primer nucleotide sequence, can substantially impact results generated from this, or any, bioinformatics analysis. While MSH2 and MSH6 were included in the GSE113513 identification dataset, their T-scores were insufficient to make the 500 gene cut-off required of GSEA to maintain statistical accuracy. However, this is an inherent limitation with the usage of GSEA, based approach which can only be overcomed by switching to a non-GSEA based approach.However, if the desired outcome is a prioritized list of potential gene candidates for further laboratory or clinical examination, the GSEA-based approach used here suffices.

A lack of direct experimental or clinical evidence substantially limited the conclusions drawn from this study of purely bioinformatics comprised work. Follow-up experiments must be done using laboratory techniques, such as Western blotting or qRT-PCR, to confirm top gene candidate predictions would support these conclusions drawn exclusively from mRNA expression data using GSEA based approach.Further analysis examining gene expression data from colonic biopsy samples undergoing any type of treatment for CRC that mimics clinical samples is needed to assess the prediction of results potrayed here. Further, gene expression data directly from CRC patients would be of particular interest to further explore the results generated here. Such an examination of gene expression data from treated and untreated human biopsy colonic CRC tissue samples would be limited due to challenges acquiring samples from tumor location.

## 5. CONCLUSION

This work used mRNA micro-array expression data to predict genes potentially involved in developing CRC tumours examining gene signatures. Through a GSEA-based meta-analysis approach, 54 over-expressed genes, most important ones being SLC25A32, SLC22A3, CA12, CA4, CA1, and SLC4A4, were identified amongst the 54 over-expressed genes as being most associated with CRC tumorogenesis regardless of it’s association with any other glandular tumour in origin. Overall, this work demonstrated the usefulness of a meta-analysis approach used previously to detect genes associated with SARS infection in identifying genes associated with DIPG treatment through application on mRNA expression data.Also, further laboratory, wet lab experiments and clinical examination into the role these identified gene expression changes play in developing CRC tumours might improve treatment options available and life expectancy for CRC patients.

## 6. ACKNOWLEDGEMENTS

Thanks to Dipak Vaja for graphical assistance and helping out with visualization analysis.

